# Large Language Models Can Extract Metadata for Annotation of Human Neuroimaging Publications

**DOI:** 10.1101/2025.05.13.653828

**Authors:** Matthew D. Turner, Abhishek Appaji, Nibras Ar Rakib, Pedram Golnari, Arcot K. Rajasekar, K V Anitha Rathnam, Satya S. Sahoo, Yue Wang, Lei Wang, Jessica A. Turner

**Author notes:** **Correspondence:** Matthew Turner.

## Abstract

We show that recent (mid-to-late 2024) commercial large language models (LLMs) are capable of good quality metadata extraction and annotation with very little work on the part of investigators for several exemplar real-world annotation tasks in the neuroimaging literature. We investigated the GPT-4o LLM from OpenAI which performed comparably with several groups of specially trained and supervised human annotators. The LLM achieves similar performance to humans, between 0.91 and 0.97 on zero-shot prompts without feedback to the LLM. Reviewing the disagreements between LLM and gold standard human annotations we note that actual LLM errors are comparable to human errors in most cases, and in many cases these disagreements are not errors. Based on the specific types of annotations we tested, with exceptionally reviewed gold-standard correct values, the LLM performance is usable for metadata annotation at scale. We encourage other research groups to develop and make available more specialized “micro-benchmarks,” like the ones we provide here, for testing both LLMs, and more complex agent systems annotation performance in real-world metadata annotation tasks.

## 1 Introduction

Scientific publications are highly stylized writings with strong rules and norms for formatting and arranging the information that they convey, unlike the freedom afforded to writings like web pages, business letters, and other working literature. This regularity allows scientifically literate readers to deftly read and process these publications. Despite these restrictions, scientific publications are generally free-text documents, and the freedom afforded to their writing allows an incredibly high degree of variability which limits the discovery of relevant publications within the broader research workflow. Basic text search (e.g. Google web search, PubMed, etc.) has often been ineffective as it has worked, historically, at the word and phrase (syntactic) level, which detaches the search terms from much of the surrounding (semantic) context. For example, a web search or searching a collection of documents for a term, such as “schizophrenia,” will turn up many papers that mention schizophrenia in passing or even in the context of “without schizophrenia”, and those papers limit the findability of papers which present relevant research results directly relevant to the topic.

Historically, human curation of the scientific literature has been required to overcome these limitations, but this approach fails as the scale of scientific publication increases (Tan et al., 2024; Yadav et al., 2024). *Curation* refers to the extraction or development of specific, context-relevant, information from publications and the organization of this information into “annotations” which we treat as metadata that are attached to the original publications (Tan et al., 2024). This metadata specifically allows programmatic methods for accessing and further processing this published literature. This “further processing” may include finding links to data attached to the publications, as well as processing the literature for systematic or quantitative reviews.

This literature curation and metadata annotation comes with a number of challenges. Perhaps the foremost of these is economics: outside of a few projects such as PubMed or corporate databases, there are very few resources available for detailed human curation of the scientific literature. But even if such resources existed, there are still problems. First, the task requires expertise in the relevant scientific area or specialized training of the annotators. Second, annotation requires substantial attention to detail, as well as high levels of focus, both of which are difficult for human annotators to achieve. Finally, the task is intrinsically monotonous which makes it difficult for people to maintain accuracy for more than a short period.

Large language models (LLMs) have disrupted artificial intelligence and machine learning research in various ways across multiple domains of use, including science (Fernandez et al., 2023; Maik Jablonka et al., 2023; Thirunavukarasu et al., 2023; Zhao et al., 2023; Nejjar et al., 2024; Sahoo et al., 2024). Within scientific research, LLMs are being used for a variety of tasks, but here we focus on their use in processing the scientific literature itself. Prior work has shown that LLMs can be used in several relevant natural language tasks such as summarization of both single and multiple documents or gathering materials for reviews (Lála et al., 2023; Agarwal et al., 2024), and annotation (Ding et al., 2023). Here we explore the capabilities of one of the most sophisticated of these models (GPT-4o from OpenAI) to do an annotation task traditionally done by human experts.

Given the impressive performance of LLMs in many tasks it seems obvious to try them on the annotation task. However, results have been mixed: Wadhwa et al. (2024) find that GPT-3 achieves state of the art performance in relation extraction from texts while Kristensen-McLachlan et al. (2023) find that LLMs are unreliable as annotators for problems in tweet classification. Aldeen et al. (2023) find that LLM performance in annotations vary widely by the specific annotation task tested. These results suggest that LLM performance on annotation will depend critically on two things: the specifics of each annotation task itself and the choice of LLM used to do the task.

When considering the capability of LLMs to do metadata annotation we must acknowledge that the human-made gold standard data, which we compare to machine annotation, is far from perfect. In a previous study (Sahoo et al., 2023), we used trained undergraduate students to do annotation of neuroimaging papers with metadata relevant to understanding the type of experimental data reported in these papers. Even with well-trained students and multiple quality control measures (using multiple annotators per annotation, more experienced students reviewing the work of junior students, multiple passes over the annotations, reviews by subject matter experts, etc.) there were many errors in the final set of annotations. Additionally, the co-developed Neurobridge ontology, like all non-probabilistic ontologies, has crisp boundaries which require arbitrary decisions in terminology. This leads to the inconsistent application of the terms when annotating.

We take the position that annotating publications with machine readable metadata extracted or generated from the unstructured text of the publications is sufficiently valuable to warrant its creation. One potential criticism of this approach could be that LLMs can interact with unstructured text directly, therefore what purpose does such metadata serve? While it is possible that LLMs could, for each new problem, process many publications directly, there are issues with this approach. First are the direct costs involved. Commercial LLMs do these sorts of tasks well, but open weight LLMs (such as the Llama series of models from Meta) do not currently perform as well, as can be seen from the various LLM leaderboards. Second, there are serious concerns about the environmental impacts, both energy and water usage, for the large commercial models (Strubell et al., 2019; Hisaharo et al., 2024; Jiang et al., 2024). If eventually open-weight LLMs could take on these tasks with less damage to the environment, there are still the compute costs and the direct work of deploying and maintaining these models. (Running an LLM, even at a moderate scale, is not as simple as downloading and running an executable.) Therefore, minimizing the number of times a publication must be processed has intrinsic value. Finally, while LLMs are capable of sophisticated processing of publications, they are not instantaneous, so passing many publications through an LLM workflow is, and will likely remain for the near-term future, unacceptably slow for on-demand use.

Preprocessing to extract such metadata and curating the results as a searchable repository for the future enables reuse and repurpose, thus saving valuable resources. Further, the benefits of having such a repository are additive as it enables aggregation with additional metadata based on other future LLM extractions. We believe that a growing prompt-annotated metadata repository would be a very useful tool for researchers who can search for past annotated data and add to the repository with focused annotation prompts that they find relevant for their current research. *Community annotated metadata using AI-prompts will be a valuable addition for future research and improve the metadata signatures for datasets with minimal effort*.

Many neuroimaging papers describe research studies where the authors make their data available, but this is often done without putting the data directly into public repositories or making the data easily findable via internet search. The goal of the Neurobridge project (Sahoo et al., 2023; Wang et al., 2023a, 2023b) is to facilitate the ability to find neuroimaging data described in the published literature. With this goal in mind, the characteristics of a magnetic resonance imaging (MRI) dataset that we have annotated here address the question: “If I received this dataset from the authors, what would I receive?” Specifically, we focus on aspects of the data that allow either data reuse or which could be used to address further questions, such as: What types of MRI scans were collected? Who were the participants who were recruited? What were the parameters used in the scanner? Therefore, we are interested in descriptions of subject or participant populations, including diagnostic status; the structure of the experimental groups; anatomical MRI imaging parameters; and classification of functional MRI methods (e.g., resting state MRI versus task-based MRI); and detailed descriptions of task-based MRI, with classification into specific categories. The Neurobridge Ontology serves as standardized terminology that can be used as annotations. This list of questions will expand as Neurobridge continues.

We present examples of three specific targeted LLM annotation tasks in this paper that relate to data findability. We note that this same information is of value in the curation of papers for systematic reviews and meta-analyses. Specifically, we develop several “micro-benchmarks” that are directly relevant to our curation problem, and we evaluate the leading large language model on this annotation task. By “micro”-benchmark we mean an exceptionally well curated, therefore usually *small*, collection of test cases with gold-standard human extracted metadata. (We might call such extremely vetted data a “platinum” standard, but there is no term in current use for such data.) We expect this collection of micro-benchmarks to expand as part of the ongoing Neurobridge research and encourage the development of additional benchmarks by other research groups using LLMs for these types of annotation tasks.

We provide the benchmarks we have developed, along with all testing materials and code for comparing model performance. All materials are available at this paper’s GitHub repository, please see the Data Availability Statement. We expect this repository (or a linked one) to eventually contain additional test sets, papers, and labels, for other annotation targets and we welcome contributions from other research groups working on similar annotation problems.

## 2 Methods

### 2.1 Experimental Data

The text used in these experiments are the published texts of scientific papers (hereafter “publications”) and several sets of human annotations treated as standards for comparison with the LLM results or used as performance comparisons. We used a subset of 186 open access full-text publications from the PubMedCentral (PMC) as our main collection of scientific papers in this study (see below). Several sets of highly curated labels, available on subsets of these publications, were used to jump start annotations as described below.

#### 2.1.1 Annotations: Targets

Human-assigned annotations were either available, partially available, or newly produced for the following aspects of the publications. The previously available annotations for these tasks are all taken from (2023).

Each of these sets of annotations defines a specific annotation task:

- **Task 1: General Imaging Type** - (25 publications) The 3 labels available for this annotation are broad categories of types of human neuroimaging: (1) T1 weighted anatomical images, (2) resting state functional MRI, and (3) task based functional MRI. This is the simplest category of labels we used and only requires classifying the general neuroimaging modalities reported in the publications.
- **Task 2: Structural Imaging Parameters** - (44 publications) These are the parameters used to collect T1 weighted images, such as magnetic field strength, TR, TE, etc. (See table 1 for a full listing.) There are 12 of these parameters we identified as being the most used to describe anatomical imaging, but it is common to just give a subset of these 12, as some of the parameters can be derived from combinations of the others. Which parameters are explicitly stated in a publication is a choice by the authors and, while guidelines exist, individual presentations can be idiosyncratic. While more complex than general imaging type, this task requires finding the specific parameters that are listed in the publication and recording only those explicitly present. These annotations were prepared expressly for this research and full details of how they were made are given in the next section.
- **Task 3: Experimental Group Information** - (30 publications) From previous research we have trained annotator labels for the participant (subject) groups used in the research presented in each of the 30 publications. Specifically, there is a set of 41 diagnostic labels as potential annotations and the annotator identified which label is appropriate for each experimental group presented in the publication. Here we require the LLM annotator to determine the number of participants in each group as well, which the human annotators were not asked to do previously. These participant count annotations are new to this work.

These three annotation tasks present a variety of challenges for the annotation of neuroscientific publications.

Task 1, the *general imaging type* is the simplest task as it only requires recognizing that something is present in the text, usually presented in clear or highly regularized language, and only requires assigning a generic label to indicate that the thing’s category is present. As such it constitutes a classical multilabel classification problem (Tsoumakas et al., 2006).

Task 2, identifying the *structural imaging parameters* is also a simple annotation case, only requiring copying values present in the text. Although the parameters for T1 structural images need to be stated explicitly in the methods of each paper, there has been a lack of standardization in how these parameters are reported, and some of the parameters can be derived from others. This creates a complex system of reporting where there are many different combinations of reported parameters that may imply the same underlying set of scanning parameters (see below).

Additionally, this requires a sensible grouping of the parameters by scan, as some parameters are present but have different values in different scans (T1, T2, etc.). For the annotations to be correct, the parameters reported must belong to the correct scan. While this is generally easy for humans, machines may confuse or combine parameters from different scans. Our pilot studies of this process on earlier OpenAI models such as GPT-3.5 and GPT-4 showed many errors of this type. We also note that our initial human annotations contained significant errors as well (discussed below), so again we emphasize that human annotations are never perfect without intense review, revision, and curation.

We selected 12 structural imaging parameters for extraction by the LLM (see table 1). Every paper should report a subset of these parameters. We expect the LLM to simply copy the values if present in the paper and report the value as is, with one exception. That exception was “scan acquisition time” which, when present, we requested to be reported in units of seconds no matter how it was listed in the source publications. (Acquisition time was reported in the original publications in a wide variety of different formats and units.) There are some minor differences in what was annotated by the student annotators and the LLMs, see the discussion of the annotation process (below) for more details.

**Table 1.**
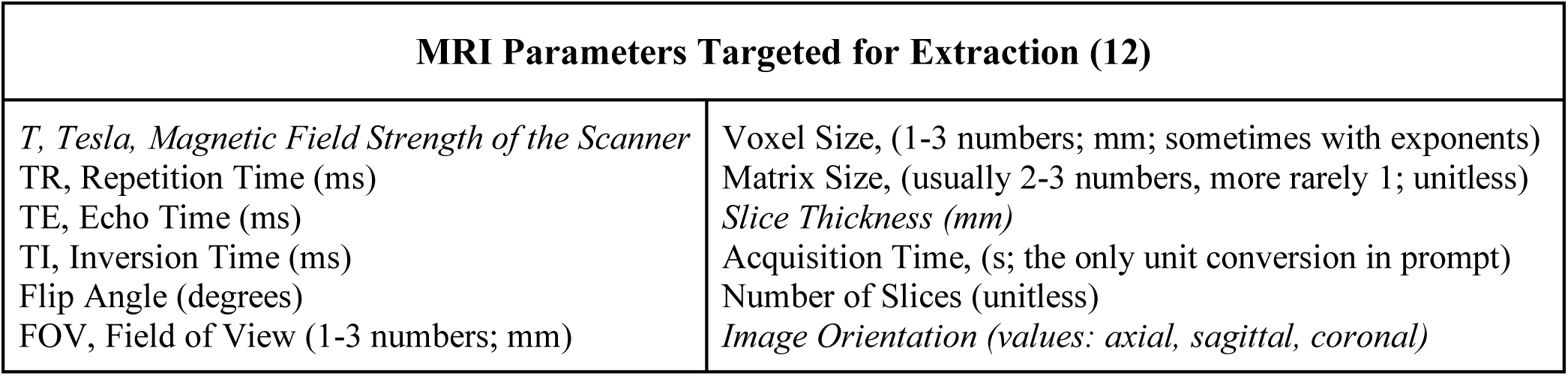
The structural MRI parameters targeted for annotation from each publication (task 2). Graduate student authors labeled the annotations in Roman font, and a senior author labeled the items in *italics*. See the discussion in the text for details.

| MRI Parameters Targeted for Extraction (12) |  |
| --- | --- |
| *T, Tesla, Magnetic Field Strength of the Scanner*<br>TR, Repetition Time (ms)<br>TE, Echo Time (ms)<br>TI, Inversion Time (ms)<br>Flip Angle (degrees)<br>FOV, Field of View (1-3 numbers; mm) | Voxel Size, (1-3 numbers; mm; sometimes with exponents)<br>Matrix Size, (usually 2-3 numbers, more rarely 1; unitless)<br>*Slice Thickness (mm)*<br>Acquisition Time, (s; the only unit conversion in prompt)<br>Number of Slices (unitless)<br>*Image Orientation (values: axial, sagittal, coronal)* |

Task 3, the *experimental group information* is the most complicated of our annotation tasks. Here we chose a set of publications where each publication had exactly one or two participant groups. The process requires that an annotator do the following:

1. Read the language used in the publication to describe the experimental groups and determine the psychiatric diagnoses that are present in natural language. Note that this language is not fully standardized across either area of research or publications. As one example of a standardization issue, we note that this language has changed over time: with the publication of the Diagnostic Standards Manual 5 (DSM-5) the separate diagnostic labels “alcohol dependence” and “alcohol abuse” were combined into “alcohol use disorder.” So, as an example, the distribution of terminology is nonstationary.
2. The description in the publication must be mapped into the set of terms in our controlled vocabulary. Even assuming perfect reading of the publications and full understanding of the controlled vocabulary, it is possible for annotators to choose different terms given intrinsic vagueness of language.
3. The label for the group must be paired with the correct final count of subjects or participants. Any cases that might have the correct label, but which lack the correct count, are considered errors.

These operations are usually easy for people but have only recently been consistently achieved by LLMs. In our pilot studies, the GPT-3.5 and the first version of the GPT-4 models could not consistently do this task, often inventing new terms that were not in the controlled vocabulary (“hallucinating” in LLM terminology). Note also that in Sahoo et al. (2023), the students were not required to annotate the group sizes so we do not have a human comparison for these.

#### 2.1.2 Annotations: Process

##### 2.1.2.1 Previous Work: Annotation Tasks 1 and 3

Tasks 1 and 3, General imaging type and the experimental group information, were annotations from previous work. Briefly, both sets of annotations were made by trained (undergraduate) student annotators in several passes over the publications. During the first pass, students entered their choices into a spreadsheet and, when relevant, made recommendations for new ontology terms to the ontologists; the Neurobridge ontology was co-developed with the annotations. After consultations between the students and the ontologists, the draft ontology labels were entered into the spreadsheets. In a later phase, the raw text of the publications was marked up directly with the annotations in specially formatted files by a new group of trained students. During this latter phase the earlier work was checked by the new students and, finally, reviewed by “senior annotators” (students who were promoted from the annotation task or their supervisors) for correctness. See Sahoo et al. (2023) for full details.

As noted above, this process did not produce perfectly correct annotations. In fact, despite the *extensive* effort put into the process, many errors were present in the final collection of 186 publications annotated.

In this study, the authors MDT and JAT reviewed, evaluated, and corrected these student annotations that were used in this study, producing new much more extensively reviewed and corrected set of annotation labels for these two tasks. These were vetted repeatedly, and the final annotations were only accepted when both authors agreed. Additionally, after the LLM annotation, each of the disagreements between the humans and the LLM were reviewed again. This process was repeated until every disagreement between humans and machine was accounted for; this yielded the final gold standard we now make available in this work. We note that this final set of gold standard annotations has been subjected to more extensive review than most “gold standard” evaluation data.

##### 2.1.2.2 New Work: Annotation Task 2 – Structural Imaging Parameters

The structural imaging parameter annotations were made by graduate student authors (NAR and PG) and reviewed by senior authors (MDT, JAT). The graduate students were required to label the set of 44 publications for 9 possible values (see table 1). So, there were 9×44 = 396 annotation positions (potential parameter values) to fill in. Note that three of these labels were for FOV, matrix, and voxel size. Each of these parameters may be represented by 1, 2, or 3 numbers. The students reported the values as strings, not as individual numbers, so each of these count as a single annotation position.

The LLM treated these 3 items as 9 distinct annotations, one for each number reported. This will affect the count of potential annotation positions in the discussion below. The graduate students each labeled about half of the publications annotated, then they switched and reviewed each other’s work. Below we call this set of annotations (before review) the “graduate student annotations.”

These annotations were reviewed and normalized by author MDT. Normalization corrected for different encodings: some of the elements annotated by the students were cut-and-pasted while others typed, so different symbols, sometimes visually indistinguishable, were all corrected to specific standard symbols. Additionally, units were normalized (e.g. “10 degrees” and “10°” were both made the same) and generally removed from the student annotations and embedded in the format used to store the data (that is, either paired with a column name in spreadsheets or stored as a different data element in JSON formats, see task 2 below for the JSON). OpenRefine (version 3.8.2) was used as the primary tool for this cleaning process (Ham, 2013; Petrova-Antonova and Tancheva, 2020). We note that during this review and normalization process author MDT added the three remaining annotations: slice thickness, image orientation, and Tesla (magnetic field strength). The annotations added by author MDT are referred to as the “additional annotations” below. This led to an additional set of 132 (44×3) annotation positions.

Subsequently, all the annotations were reviewed by both MDT and JAT. Note that by the end of this, some annotations have been through 5 passes: original annotator, second annotator, MDT (during normalization), JAT, then MDT again, while the ones added by MDT have been through 3 passes (annotation by MDT, followed by review from JAT and MDT). We call all of these final, thoroughly reviewed annotations the “full human system annotations.” Throughout this process, data was collected on human errors made in annotation at each stage; these will be discussed below. Note that we have three relevant stages to assess the accuracy of the annotations: the correctness of the graduate student annotations (before review) together with the correctness of the additional annotations (by MDT) before review, the annotations after being reviewed and updated by experts (JAT and MDT), and the correctness of this reviewed data during the post LLM review of differences. (Each of these can be considered a different stage of the development of the annotations.) The graduate student and additional MDT annotations are comparable to what most papers call a “gold standard” data set, despite still containing many errors. The full human system annotations are much more deeply reviewed than is usual in this kind of work. Our new “gold standard” annotations are the final set that has been through all these processes.

This particularly intense review process yielded a set of annotations that are more correct than we have come to expect from most human annotated gold standard data sets. However, after LLM annotation, the entire set of human/LLM disagreements was reviewed again, which yielded 2 additional human errors that had made it through the entire process described above. We discuss the correctness of these sets of annotations in the appropriate section of the results below and we consider the costs involved there.

All of the final gold standard annotations are available in the GitHub repository for this project, see the data availability section.

#### 2.1.3 Publication Texts Details

LLM usage. The raw text of the publications given to the LLM in two of our experiments (tasks 2 and 3) were the BioC texts of the publications (Comeau et al., 2019). These BioC formatted papers were further processed to provide the plain text of the papers with footnotes, citations, and other technical apparatus removed. Due to context window size limitations of earlier LLMs, only the content of the papers to the end of the methods sections were provided to the LLMs. This included: titles, abstracts, introductions, and methods sections; removing the results, discussion, and other latter parts of the papers. The sections were determined from the metadata provided in the BioC format. For task 1, general imaging type, the LLM was given the PMC provided PDFs rather than the BioC text. This task’s workflow was dependent on the LLM provider’s internal process for converting the PDFs into usable text for the LLM and, as such, we do not have access to the details of how this was done. We note that there were two runs which generated failures in parsing the PDFs. This appeared to be an intermittent fault and in both cases repeating the analysis in a new chat session resolved the problem with no changes. See below for more details (section 2.2.1).

Human usage. For annotation tasks 1 and 3, the human annotators had access to the full (all text sections) BioC text for their final markup, but they were not restricted to using only this text and were free to review PDF versions of the papers. For the structural imaging parameters task, the human annotators used the PMC provided PDFs as their texts.

### 2.2 Large Language Model and Prompting Strategies

#### 2.2.1 LLM: GPT-4o

The present study focuses exclusively on the GPT-4o (omni) LLM, the 2024 flagship product from OpenAI, although we developed many of the experimental tasks on earlier models (GPT-3.5 and GPT-4 at various checkpoints). It is used as the reference model as it is an extremely capable model with a reasonable price point for use in research applications. As the goal of this paper is to establish whether the most powerful models available can reasonably solve these tasks, we did not attempt to systematically explore all currently available models but instead established a baseline against one of the most prominent current commercial models. As there are new models released continuously, overall LLM model performance is a rapidly moving target. We provide code in the paper’s GitHub repository that can be modified to enable testing other LLMs with our annotations.

We used two methods of submitting our prompts to the LLM. Task 1 was run manually via ChatGPT (OpenAI’s chat interface), selecting the GPT-4o model and uploading the PMC-provided PDFs of the publications for analysis. Each paper was run with the prompt in a fresh chat session, using the default parameters for ChatGPT. Accessing GPT-4o via the chat interface precludes knowing the specific model version or “checkpoint” used; however all of these were run on the same day in August 2024, so they should all have used the same version of the model. The other tasks (2 and 3) were run through OpenAI’s API, which allows direct programmatic access to the models via functions in a corresponding Python library released by the company (github.com/openai/openai-python). The model used in those tasks was the GPT-4o LLM (version checkpoint: gpt-4o-2024-08-06). In these experiments, the raw BioC text of each paper in turn was appended onto the end of the prompt for the task and submitted for processing. As LLMs are stateless, this approach is the equivalent of using a fresh chat session as in the manually run experiment.

For these tasks, all LLM parameters were set to their defaults except for the “temperature” parameter. Temperature appears to be related to LLM response variability, although this claim is contested (Renze and Guven, 2024). Others commonly claim that it is a response “creativity” parameter, but this is also unlikely (Peeperkorn et al., 2024). For task 1 the temperature was left at the default for the ChatGPT interface. For task 3, the temperature was set to 0.2 (low), based on informal advice we received, but we did not explore other settings. For task 2 we had to set the temperature to zero to fix a specific problem. This task used the most complex JSON prototype of all the tasks. When the temperature was greater than zero, the LLM would often return simplified JSON, that is, it would pick out the annotations present in the publication and return only the JSON fields corresponding to the parameters explicitly reported. The JSON fields that corresponded to parameters not present would often be elided in unusual ways that were problematic for post-processing. Setting the temperature to zero made the LLM always return the exact same JSON each time, just with the JSON nulls replaced with values for any parameters present in the publication. All the other parameters remained present in the JSON and set to null. This is the behavior that we wanted. For the other tasks, no temperature changes were needed to maintain the expected JSON format. We take the position that JSON should always remain fully present (i.e., don’t delete fields for information not available) to simplify the postprocessing of the JSON returned from the LLMs.

#### 2.2.2 Prompting Strategy

Given that we were using a capable leading edge LLM, we explored the quality of annotations under only the simplest possible usage (Schulhoff et al., 2025). Specifically, we used “zero-shot” prompts for these tasks (Reynolds and McDonell, 2021; Li, 2023). Prompts are the text that specify the task that the LLM is supposed to perform (Bhandari, 2024). A zero-shot prompt is one in which instructions for the task are given, but no specific examples are included. Performance with a zero-shot prompt can be thought of as a type of baseline performance or naive prompting strategy. If there are annotations that do not perform well under zero-shot prompts, so-called “few-shot” prompts can be used to improve performance. These prompts include both the task instructions along with one or more examples. If there are known cases that are challenging for the LLM, these problems can often be solved by adding examples that specifically cover the most challenging cases. For more complex annotation situations there are sophisticated systems, such as DSPy (Khattab et al., 2023a, 2023b), that exist to automate few-shot prompt development with examples.

We selected the zero-shot approach for several reasons, the main one being that this approach is what most people would try first when faced with a new annotation problem. Current commercial LLMs are quite capable when used with zero-shot prompts, and any annotation problem which requires going beyond simple prompting strategies will likely require addressing specifics that may be unique to the class of annotations involved. Perhaps obviously these procedures are not perfect. However, human annotation has its own problems as noted above. No method, either human or automated, will ever annotate perfectly except in situations of extensive review and curation, but we find that the types of errors GPT-4o commits under this strategy are comparable to human annotation (particularly when that human annotation is given reasonable review and verification).

Additionally, our prompts contained JSON prototypes of varying complexity that guided the LLM responses in each annotation problem. See figure 3 for the most complex example. We recommend using more detailed JSON, like that for task 2 here, as such prototypes contain information that improve LLM performance in annotating publications. The handling of the JSON format by LLMs has dramatically improved over the course of this study. The most recent OpenAI models offer a new feature, called ‘structured outputs,’ that allow for more fine-grained control of the JSON returned from the LLM (platform.openai.com/docs/guides/structured-outputs). We did not use these new features as they became available too late in the course of this study. All the outputs from the models presented here used only the more basic JSON-mode provided by OpenAI for their models along with in-prompt instructions to return JSON as the response (platform.openai.com/docs/guides/structured-outputs/json-mode#json-mode). Despite using this older and more limited control over the LLM output, we still achieved excellent results.

#### 2.2.3 Evaluation

The tasks described here are evaluated in two important ways. First, the tasks are analyzed using simple accuracy (the ratio of correct annotations, including “not present” or “not applicable” as required, to the count of all possible annotation positions). However, given the high accuracy reported here (all greater than 90%), we focus on comparing these numbers with human performance for calibration. Second, for each task, there is a qualitative analysis which reviews each of the disagreements between the LLM and the human gold-standard data. It is known that LLMs occasionally do better at annotating than humans do (Nahum et al., 2024). We also find this in our results. This includes both finding additional errors made by humans that slipped through review and occasionally finding that the LLM annotations are better organized than the human work. Note that with this review we were compelled to change our conception of our gold standard, producing a higher quality final product than previous human-centered annotation processes have produced.

## 3 Results and Discussion

### 3.1 Annotation Task 1: General Imaging Type

This task used the prompt given in figure 1. An issue arose in this case: for nine of the papers the given prompt failed to generate correct answers. For these papers, a new run was started and a prompt that said, in its entirety: “Please summarize this paper” was given. Then, after ChatGPT generated the summary of the paper, the original prompt was repeated verbatim; and this action substantially improved the performance of ChatGPT. Note that this is a chain; these two prompts were run in sequence in one session. This action improved the results even though the summaries were not relevant to the errors present in the initial response. Adding this summary prompt before the papers that worked initially did not change their annotations, so a fully automated process for this task via the API is trivial to implement.

**Figure 1:**
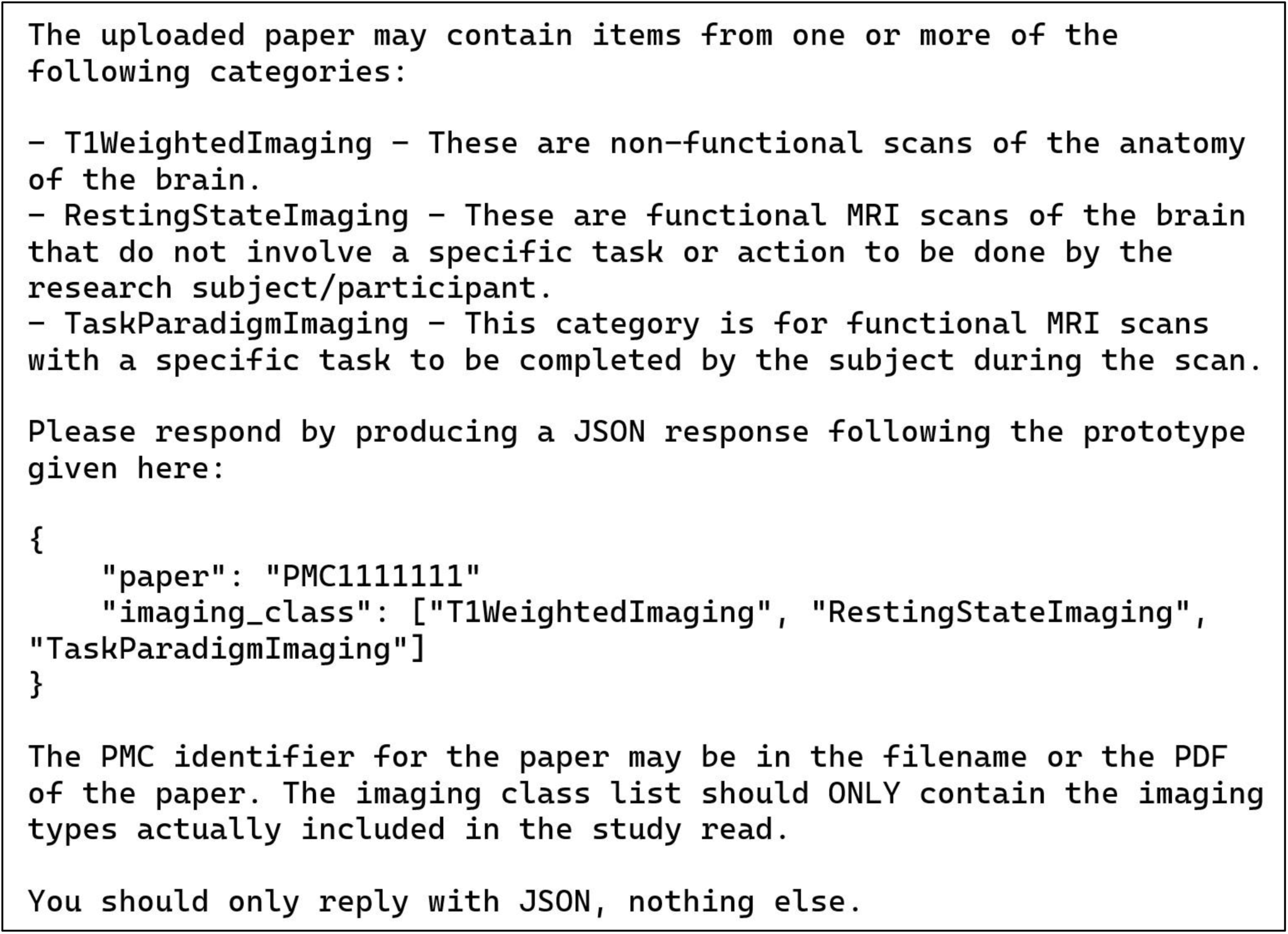
Manual zero-shot prompt for general imaging type task (task 1). This text was pasted into the chat window after uploading the publication PDF. See text for details.

The scores for this manual process are presented in table 2. We scored the LLM based on both the initial response to the manual prompt and again with this prompt being repeated after generating a summary. The initial LLM responses for the publications analyzed are presented in the column “LLM without summary.” Simple accuracy here is only 84% reflecting 12 errors: 8 omissions (all “T1 Weighted Imaging”) and 4 false positives (all “Resting State Imaging”). After summarization, this was reduced to two errors, both false positives for “Resting State Imaging.” Both the LLM with summaries and human student annotators scored an accuracy of 97.3% on 2 errors for each (see below for details of the errors). It is reasonable to expect these labels to be among the most easily assigned, so this result with LLM and humans tied is not surprising. It also makes the point that human annotations are never perfect. The second row of the table is the fraction of “perfect” cases out of the total number of cases.

**Table 2.** Results for the general imaging type annotation task (task 1). The LLM was tested under two conditions: once with just the basic prompt and a second time with this prompt following a request for the LLM to summarize the publication. See text for details.

| Parameter | *LLM without*<br>*Summary* | *LLM with Summary* | *Student Annotators* |
| --- | --- | --- | --- |
| Accuracy | 84.0% | 97.3% | 97.3% |
| Fraction of Perfect Labeling | 14/25 | 23/25 | 23/25 |

#### 3.1.1 Qualitative Analysis

The student annotators are scored as committing two errors, one false positive and one false negative for T1 Weighted Imaging. One of these is correct but based on information not included in the actual publication analyzed (the information was included in a supplement to the publication). As such it is unfair to score the students as wrong here (however in the ground truth this was set to “no T1 Weighted Imaging” to accurately reflect the contents of the publication text). The other error that the human annotators made was missing an oblique reference to T1 imaging in one paper. However, it is worth noting that this paper did not follow standard reporting practices, so the reference was easy to miss, although the LLM did both find this obscure reference and correctly interpret it.

The LLM (with summarization) committed two errors as well, both of which were the inclusion of the “Resting State Imaging” indicator when this type of imaging was not present (false positives). Without summarization beforehand, the LLM made this error four times, summaries resolved two of these. In the non-summary condition, the LLM also made eight errors (false negatives) for the T1WeightedImaging label. Given the prior probability of structural imaging being included in almost any given fMRI study this seems strange. Also, as noted, none of the generated summaries included any mention of structural imaging, so it is not clear why these summaries improved the performance in so many cases.

### 3.2 Annotation Task 2: Structural MRI Parameters

The prompt and the JSON prototype for this task are presented in figures 2 and 3. Note that these are shown separately for the presentation here due to their size but were combined with each other and with the publication text to be analyzed into a single block of text when sent to the LLM via the API. The instructions in figure 2 simply list the parameters to be collected, along with a one-line definition of what the parameters mean, and an admonition to only list parameters explicitly present in the publication. This example uses a more complex JSON format than our other tasks in that it also includes embedded information to guide the LLM toward a correct solution and alignment with the instructions. Each of the items shown in figure 3 includes a brief informative description and the most common units of measurement used for each parameter. Although occasionally the publication text strays from these units, the LLM successfully mapped the parameters in the free text into the JSON with units used correctly. We did not ask the LLM to do any unit conversions, except for the “acquisition time” of the scans, which was reported in a wide variety of ways in the original publications. We prompted the LLM to convert any different time representations into seconds no matter how they were reported (figure 2). This is the only explicit demand for unit conversion. Amazingly, given the wide variety of ways that acquisition time was written in the publications, the LLM was successful in converting and recording these for all but one case (see below).

**Figure 2:**
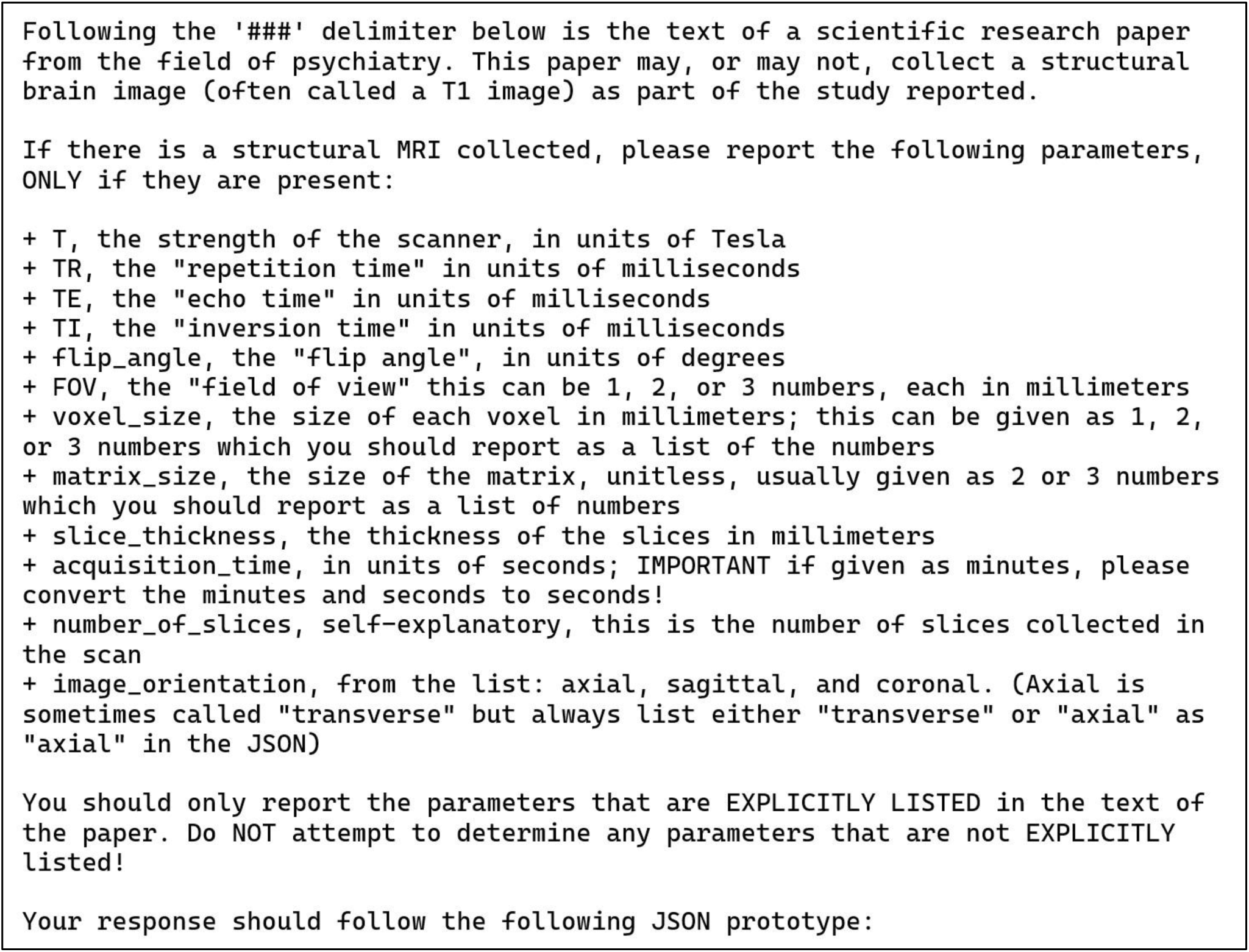
Zero-shot prompt for the structural MRI parameters task (task 2). This prompt describes the potential MRI parameters to be returned. It provides a limited vocabulary for “image orientation” and the expected units for all other parameters. Also, it tells the LLM to convert scan acquisition times to units of seconds, no matter how this was originally reported in the publication.

**Figure 3:**
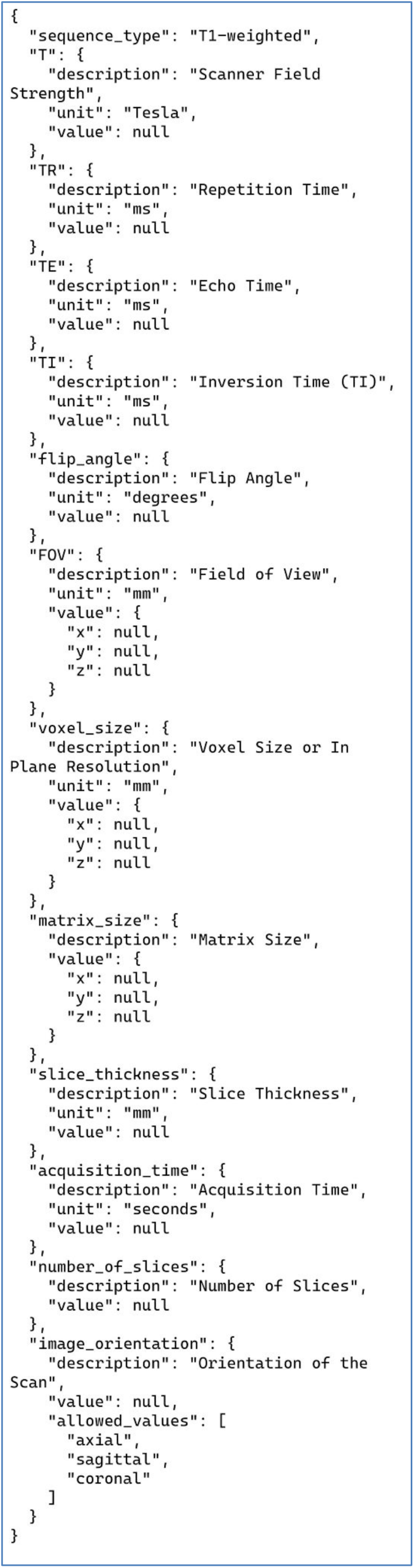
JSON prototype for the structural MRI parameters task (task 2). Each entry corresponds to one T1-weighted structural MRI parameter, and includes the standard units used for that parameter to guide the LLM response. The LLM returns this JSON in its entirety with any parameters present in the text replacing the relevant “null” values under the “value” keys. Parameters not explicitly listed in the publication text are left as JSON nulls. Note the use of allowed values for “image orientation.”

Out of 792 annotation positions in the JSON (44 publications × 18 possible values) the LLM annotations did not match the gold standard annotations 35 times, for an overall (preliminary) accuracy of 95.6% *under the assumption that the gold standard is fully correct*. However, not all mismatches will turn out to be wrong, see the next section.

We reiterate that most of the values to be filled in are nulls, however we consider missing a value that is present to be as problematic as filling in a null that is not present. We also note that both the LLM and the humans made both types of errors.

#### 3.2.1 Qualitative Analysis

As the number of mismatches was small, we reviewed all 35 of them to determine the differences between human and LLM annotation. We include the PMC ID numbers and citations for any specific papers we review in detail.

The first group of mismatches are differences from the gold standard that are, in fact, correct. In one paper, PMC5037039 (Janes et al., 2016), we discovered that the LLM reported the Tesla rating of the scanner as 2.89T, not 3T as reported by our human annotators. In fact, both values were listed in the paper, with the 2.89 being more precise, so the response from the LLM is correct. In the JSON prototype we require 1, 2, or 3 values for voxel size. We note that the human annotators just copied these numbers from the publications verbatim (as strings) without much consideration (this is what they were told to do). The LLM recognized that papers reporting “1 mm^3^ isotropic” for voxel size corresponded to 3D voxels with values of (1 mm, 1 mm, 1 mm), and returned the unpacked x, y, z values appropriately. This may be considered an error or not, depending on the interpretation or the goals of the annotation process, but this is a correct interpretation of the notation. This specific difference occurs in 8 papers and accounts for a total of 16 of the differences with the gold standard. Combining these with the previous difference for Tesla rating, changes the mismatches from 35 to 18 yielding a revised accuracy of 97.7%, *if we prefer these changes to our original annotation scheme*. The annotation supervisors were impressed with the LLM’s recognition and correct use of the “isotropic” notation and noted that it was a poor choice in the design of the human annotation process not to require a similar unpacking from the student annotators.

The second group of mismatches are related to errors or ambiguities in the original papers. In one paper, PMC6031869 (Hua et al., 2018), the authors list the “in plane resolution” or voxel size as “231×232” which our expert considered most likely to be the values for the matrix. This is what the LLM assigned those values to; the student annotators did not. In PMC6551253 (Chumin et al., 2019), the text contains the odd expression “…Field of view = 192×168 matrix…” which the LLM labeled as FOV, but which our expert believes should be the matrix value. However, without expertise this is a hard case to resolve, although here our student annotators did choose the expert’s preferred solution. Paper PMC6677917 (Lottman et al., 2019) stated “base resolution = 256” which our expert determined implies a matrix value of 256×256. The LLM reported this as a matrix with one value (x) of 256; the student annotators ignored this information entirely. One mismatch was due to the PDF to text conversion of one of the publications. In PMC6491039 (Sawyer et al., 2019), the value for flip angle was listed as 7 degrees, but the typesetting of the article used a superscript zero character rather than a degree sign, and our raw text processing turned this into “70,” which is what the LLM read and reported, instead of 7° as was intended. This is an error of the preprocessing of the file, not an error introduced by the LLM itself.

The final group of mismatches are definite errors on the part of the LLM. In PMC6289814 (Kim et al., 2019), the LLM took the FOV values from a different scan. In PMC6104387 (Hahn et al., 2017), the LLM copied the FOV and the matrix values from an unrelated T2* image. Finally, in PMC6491039 (Sawyer et al., 2019), the LLM missed an explicit mention of the acquisition time.

In summary, while there are clear errors present in the LLM annotations, they are rare and the general types of errors made are not substantially different from the sorts of errors made by the human annotators.

#### 3.2.2 Curator/Annotator Detailed Comparison

As described in the methods section we have detailed information available about these annotations for the following stages of the annotation process:

1. The **graduate student annotations** – These are the annotations made and reviewed by the graduate student annotators. Errors here were discovered both during review and normalization stage and during the final (pre-LLM annotation) review.
2. The **additional annotations** made by author MDT – These annotations were made during the review and normalization stage. Errors here were discovered during the final (pre-LLM annotation) review.
3. The **full human system annotations** – These are the annotations that exist at the end of the entire human process, specifically after all annotation, normalization, and review passes listed above. Errors here were discovered when reviewing the individual LLM mismatches (post-LLM annotation).

We compare the accuracy of these cases here. Results of these comparisons are summarized in table 3.

**Table 3.** Results for structural MRI parameters (task 2). Accuracy of each human annotator group, annotation by LLM alone, and the complete “human system” which includes multiple review processes. See text for details of each column. The value for LLM accuracy in parentheses is the revised score for extra annotations that were correct, but did not agree with the original gold standard data.

| Annotator Type | *LLM* | *Graduate Student*<br>*Annotations* | *Additional*<br>*Annotations* | *Full Human System*<br>*Annotations* |
| --- | --- | --- | --- | --- |
| Accuracy | 95.6% (97.7%) | 94.1% | 93.9% | 99.6% |

The **graduate student annotations** required labeling 44 publications for 9 possible values. For this set, there were 9×44 = 396 potential annotation positions to fill in. The student annotators made 23 total errors for an accuracy of 94.1%. The **additional annotations** had a similar accuracy to the graduate students: three additional annotation fields across 44 publications created a total of 3×44 = 132 positions, and there were 8 errors, an accuracy of 93.9%. We note that for this task, the basic LLM accuracy of 95.6% is superior to either the human annotators (94.1% and 93.9%). If the reasonable choices of the LLM above are accepted as correct, then the LLM performance becomes 97.7%, substantially better than the coordinated and time-consuming work of three human annotators.

In addition, two errors relating to the matrix values made it all the way through both student annotation and curator review and were only discovered during the detailed review of the LLM performance. This last represents the performance of the full human annotation system as described above and in the methods. This system achieved an accuracy score of 99.6% (2 errors in 12×44 = 528 annotation positions) with a cost of *at least* 60 total person-hours of work. LLM annotation of the same texts and annotation positions cost approximately 0.90 USD and took approximately 15 minutes. Total LLM process development time took approximately 3 hours for programming, testing, etc. This comparison is not completely fair as much of the conceptual development took place during the human labeling process and normalization, but these figures provide a reasonable estimate. It is worth noting that even if human review is used, using the LLM for the first annotation passes would still be a very significant savings.

### 3.3 Annotation Task 3: Experimental Group Information

In this task the LLM was required to determine the psychiatric diagnosis and the correct final count of research participants in each experimental group present in each publication. Performance in this task is less accurate than for the other tasks, for both human annotators and the LLM. The task has the most intrinsic ambiguity as the ontology terms used are for psychiatric disorders and these often have intrinsic variability even among expert users. We note that there was variability among the ontology experts both for the exact usage of each term and which term to use for the annotation of each publication.

Each of the 30 publications used in this task had either exactly one or exactly two experimental groups (4 and 26 publications, respectively). Furthermore, each experimental group in this collection of publications had a single diagnostic label; papers with groups of combined diagnoses were excluded. This means that the LLM was required to fill 120 annotation positions, comprising 60 pairs (diagnosis and count), with each pair describing one experimental group. For the 4 publications with a single group, the other group should be left blank by the LLM; anything other than an empty response would be an error of equal weight. For these four groups null values should be given for both diagnosis and count.

The prompt for this task is shown in figure 4. This prompt is like the previous examples: it is zero-shot and provides a JSON prototype to guide the LLM. The list of psychiatric diagnoses comes from the Neurobridge ontology, which is available in the Neurobridge GitHub repository as an OWL file. (See data availability statement for details.) Note that the figure elides the list of 41 diagnoses that were given to the LLM, but the full prompt is available in the GitHub repository as well. The text of the publication was appended to the end of the prompt after the “###” delimiter which follows OpenAI best practices (help.openai.com/en/articles/6654000-best-practices-for-prompt-engineering-with-the-openai-api). Additionally, to emphasize the instruction to choose just one item from the list, we added the phrase “Please pay special attention to this requirement, my job depends on it.” These sorts of phrases were commonly used in prompt design (Li et al., 2023), although their importance has dropped off with more advanced models.

**Figure 4:**
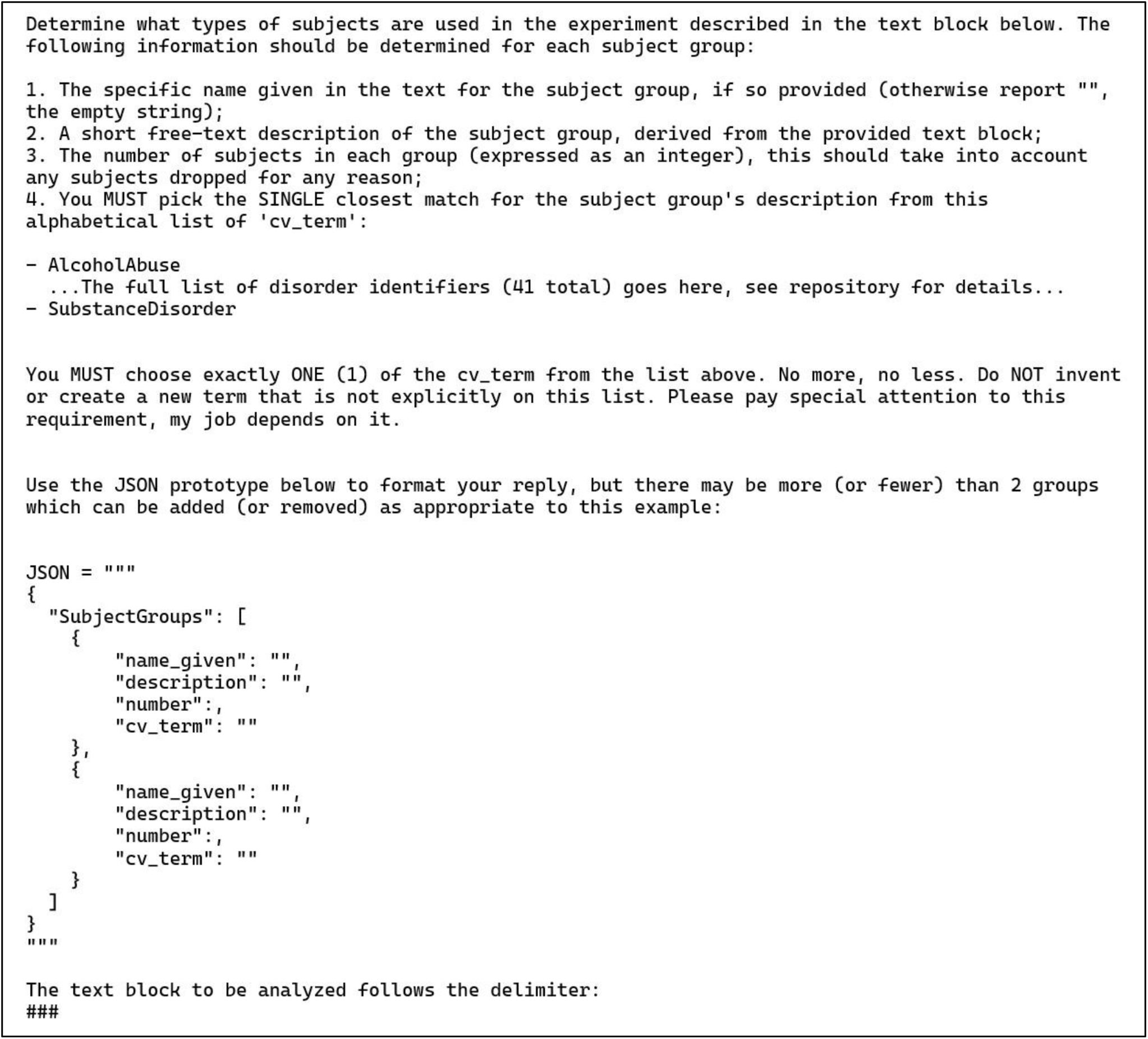
Zero-shot prompt and JSON prototype for the experimental group information task (task 3). Note that the version of this prompt shown here has had the full list of ontology terms elided. This list can be found in the GitHub repository (see data availability statement). The use of “emotional” stimuli words (“Please pay special attention to this requirement, my job depends on it.”) was a previously recommended procedure that is not used as often today. See text for details.

The accuracy of the LLM in this task is 90.8%, with 11 total mismatches across 30 publications. We note that disagreements tended to bunch together by publication: 26 of the 30 publications were labeled in perfect agreement with the human results. All errors were in just four publications, as described in section 3.3.1 (below).

There was no direct independent analysis of human annotation performance available for comparison with the 30 publications used for LLM annotation in this task. However, as part of developing the materials for this study, we reviewed 39 different publications labeled by humans as part of our prior work in Sahoo (2023). All these 39 publications were reviewed and given definitive labels by the present authors with all final decisions made by author JAT, who is a co-developer of the Neurobridge ontology. These final annotation decisions were made without knowledge of either the human or LLM annotation results. While we recognize that it is impossible to resolve all ambiguity in any ontology, this approach allowed us to develop a fully expert-labeled gold standard data set that could be compared against both previous student work and LLM performance. Our review revealed many errors remaining in the human annotation results from Sahoo et al. (2023).

Specifically, in the 39 publications reviewed that were human annotated there were 13 errors found, but this was only noted at the level of an individual publication and not at the experimental group level. That is, a publication was marked as being in error if *any* number of mistakes of annotation were made for it. Almost all these papers were two-group research studies, although a few had more groups, so the human annotators would have had to fill in labels for 78 participant groups *minimally* (2 groups × 39 publications). It was rare for multiple errors to be made in a single paper. This conservatively estimates the human annotation accuracy at approximately 83.3% (13 errors across 78 annotation positions). Also note that the collection of human annotations reviewed was not selected randomly but was based on the topics of the publications. We were specifically selecting publications connected to drug abuse which was a subset of the original collection. There is no obvious reason to suppose that our human annotators would make more or fewer errors for other topics. Table 4 summarizes this comparison.

**Table 4.**
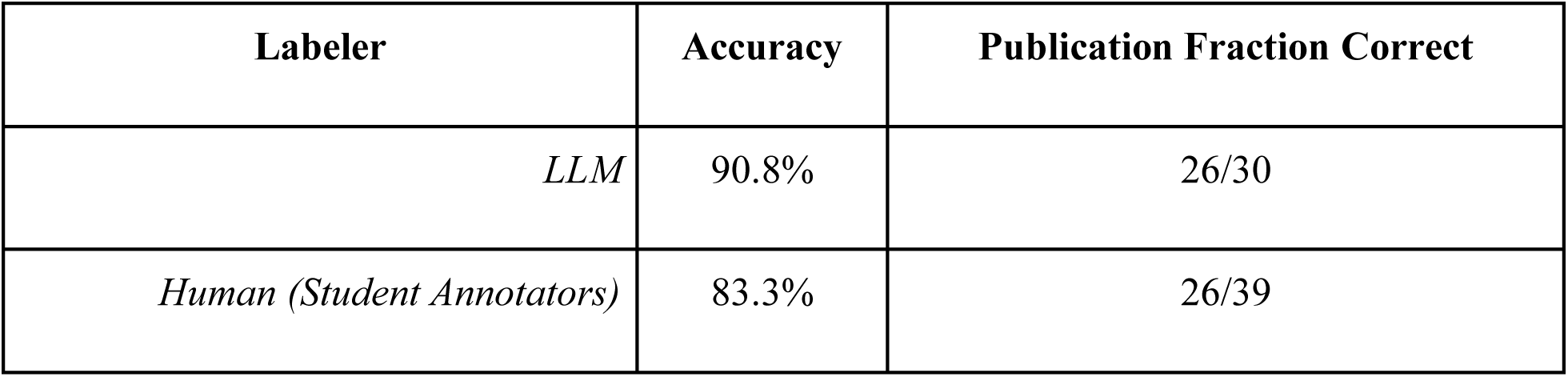
Accuracy comparison between human annotators from Sahoo et al. (2023) and the LLM annotation from this study for the experimental group information task (task 3). Note that the collections of publications annotated are different and that the human annotation performance is a conservative estimate (it is possible that it undercounts human errors). See text for details.

#### 3.3.1 Qualitative Analysis

Reviewing the mismatches between the 30-case gold standard and the LLM for this task we note the following: In the publication PMC5086261 (Forster et al., 2016), the LLM chose the annotation “substance disorder” in place of “drug abuse” and it is not clear that this difference is an actual error as these terms overlap significantly. In two cases, PMC4990879 and PMC6704377 (Chang et al., 2016; Vanes et al., 2018), the LLM appeared to ignore any control groups and split the group of interest into two, while giving each split group the same diagnostic label. We have no explanation for this strange error, as looking at the original source publication revealed no obvious reason to split these groups. In another case, PMC6215331 (Alloza et al., 2018), the LLM correctly identified one group and ignored another. The last error is hard to explain because when we took the relevant text from the paper and pasted it into a ChatGPT session (GPT-4o) with the same prompt, ChatGPT correctly identified the two groups (which happened to both be groups with “no known disorder,” or control groups). As the chat interface does not report the exact model used, this might reflect a change in the underlying model, or it may be due to the stochastic nature of LLMs.

One additional publication is also worth noting: PMC6906591 (van den Heuvel et al., 2019), although the LLM was scored as correct for this case. In this work, all the human annotators, as well as the more senior reviewers, identified two specific groups as the correct answer. This was because these two groups were the only new data collected in this publication. However, this paper has 23 total groups in it, the additional 21 being validation datasets (usually with cases and controls both, which we count as separate groups) that went unnoticed by the original human annotators. The senior reviewers noted the additional validation data but agreed that the annotators should only have listed the groups with new data, as the other data came from other publications. The LLM initially found and annotated all the groups present in the study *completely correctly*. It was scored only against the two groups the senior reviewers and original annotators had found. We were genuinely surprised that the LLM correctly annotated all these groups, especially as some of these groups were not human but primate data. We note that the prompt above does not limit the LLM to original data, so strictly this is not an error by the LLM but in the prompt it was given.

## 4 General Discussion

We describe and evaluate three real-world biomedical research annotation tasks where experimental metadata from neuroimaging research publications is collected. We show that one current flagship commercial LLM can annotate this material as well as or better than human annotators. This performance is consistent with informal AI industry assessments for other, very different, metadata annotation tasks. Even in our most challenging case, the annotation of experimental group information where the textual descriptions are highly variable and the annotations themselves contain substantial ambiguity, we achieved > 90% annotation accuracy with 87% of publications annotated perfectly according to a highly refined human standard. We note also that across these example tasks many of the LLM disagreements were not errors but instead reasonable and explainable responses.

Two main results are shown by this research. First, the real-world tasks here provide three new micro-benchmarks, as defined in the introduction, for the evaluation of systems that annotate free text with metadata that makes explicit ideas and concepts embedded in the publication text. Given the variable nature of annotation problems, assessing the performance of LLMs will require many more similar, real-world annotation tasks to fully calibrate our understanding of LLM annotation performance. Such benchmarks require much more intensive curation and review than is common in previous machine learning research as the evaluation of LLMs has shown new complications in the annotation task. Additionally, a broad collection of such applied benchmarks will be required due to the unique aspects of each annotation situation as there is no theoretical or statistical guarantee of results available for this sort of work. Our project demonstrates the first such micro-benchmarks, and we anticipate more and encourage others to make similar high-quality benchmarks available for use by the research community. An important aspect of this project is not just the results of our annotations, but the methodology identified as a way for others to follow in annotating their domain specific metadata extraction using current and future LLM systems.

The second result is that simple zero-shot prompts can get the current large commercial LLMs to do metadata extraction for cases similar to the ones illustrated here with high accuracy. We agree with informal industry assessments that large commercial LLMs are broadly capable and can perform human annotation tasks with only modest prompt engineering required. More challenging cases may require specialization of the LLM to the task at hand, either through few-shot prompting or some form of fine-tuning, which at a high level are similar processes (Khattab et al., 2023b; Soylu et al., 2024). This approach follows the principle that it is relatively simple to achieve a solution for the bulk of cases, while more effort is often needed to solve the edge or corner cases.

It is notable that the various publications do not always follow the field’s recommendations for reporting neuroimaging parameters (Nichols et al., 2016) which hinders their interpretation. This makes the annotation problem more complicated, effectively asking the LLM or human annotator to identify what the authors intended rather than what was reported. An additional challenge is the inherent ambiguity of words, even technical vocabulary. Psychiatric labels have ambiguity both due to changing diagnostic standards and conceptualization over time as well as usage that varies internationally. That is, the underlying distribution of usage of this vocabulary is not stationary either in time or space. These issues have been noted by those developing ontologies (Mugzach et al., 2015; Larsen and Hastings, 2018). Interpreting this terminology requires arbitrary choices about specific cases, deep knowledge of the underlying domain, and the ability to describe and compare nomenclature across studies.

We used JSON as our data serialization format. This choice was motivated by both the heavy standardization and broad adoption already present for JSON as an interchange format (Bray, 2017) and the fact that the output of formally correct JSON is something that all the major LLM producers have implemented. Because JSON is fully represented as text, it can be used both as an input and output format for LLMs. Specifically, JSON can be added directly to prompts while non-text formats cannot be used in this way. As noted in task 2, this allows additional information to be inserted into the overall prompt structure to assist the LLM in generating correct results. Finally, most computer languages and database systems support JSON as a format for data interchange making data extracted from publications broadly sharable. We strongly recommend the use of JSON in this context for all these reasons as there are no current standards for this sort of metadata extraction and exchange.

One concern is the presence of so-called hallucinations in LLM responses. These are outputs that contain untrue information that is apparently generated by the LLMs spontaneously. Generally, hallucinations appear when a LLM must recall some information that is embedded into the model itself rather than using information given in the prompt context. In the annotation task, LLM responses are confined to processing explicit text (the target to be annotated) and this reduces the likelihood of hallucinations dramatically. This approach is also in line with informal industry consensus. It can be hard to distinguish hallucinations from other types of errors, but in the work reported here hallucinations have played little role since, at least, the GPT-4 era.

Hallucinations must be defined based on applications. We define an LLM as having hallucinated only when it responds with something that does not exist (in the context of the task). We did find hallucinations of this sort in older models. For instance, those models would occasionally make up psychiatric diagnosis terms that had the form of our ontology’s terms (written in camel case with a similar choice of words to our terms), but which were *not* in the ontology. The current GPT-4o models did not exhibit this behavior. We do not count instances of false positives in our analysis as hallucinations. For instance, the two cases of resting state imaging labels assigned by the LLM in task 1 are not hallucinations in our sense of the term because resting state imaging is a real type of imaging that was included in our terminology for that task. Others may define hallucinations differently based on their problem domain. We also note that advances have been made in enabling LLMs to “cite” their sources so one can check the responses for hallucinations in contexts where that would make sense (Gao et al., 2023; Byun et al., 2024; Huang and Chang, 2024; Wu et al., 2024).

The larger goal that drives this work is to convert the information in a neuroimaging publication into a programmatically accessible format to enable searching the literature quickly for potential MRI datasets. The work presented here represents the first steps taken towards the goal. Additional work is required to apply this more broadly to the neuroimaging research literature. For example, we focused on human MRI experimental papers, which reported datasets that might be reasonably expected to be available for additional research use. Large-scale data aggregation studies (e.g. Thompson et al., 2020; Ching et al., 2024; Ganesan et al., 2024) entail identifying the existing datasets which might be suitable for a given meta-or mega-analysis, which requires understanding the details of each study design (Turner, 2014). Our goal is to find data that can be used to support answering new questions, therefore the analysis originally performed, the specific results of the study, and the interpretations the original authors make of their experiment and analyses are not necessarily relevant to our goals. Determining the most useful aspects of publications to annotate is an ongoing research question.

We also limited the papers to certain formats. For instance, in this work we used exclusion criteria to remove publications where the information about MRI was only located in tables. We did not provide the LLM with information from the tables, figure legends, or any supplemental material. Expanding this work to cover these cases is still needed. Additionally, automated methods are needed to verify the work of the LLMs so that these processes may be expanded to work at a larger scale. This last will likely require the use of the newest “chain of thought” models. These are all additional topics for future research.

## 5 Author Contributions

Authors MDT and JAT wrote the manuscript with significant contributions from AKR and LW. MDT, JAT, AKR, SSS, YW all contributed to the design of the study and to the preliminary data collection, analyses, and discussion. MDT did the coding for LLM processing and final scoring and designed the prompts (in consultation with JAT). Authors NAR and PG did the human (graduate student) annotations for task 2, under the supervision of SSS and AKR. Author AR did the preliminary data analyses for the tasks under the supervision of AA. All authors read and approved the final manuscript.

## 7 Acknowledgments

Preliminary analyses and portions of this work were presented at the INCF Assembly 2024.

## 8 Data Availability Statement

The materials used in this study, including the Python scripts to access the LLM via OpenAI’s API and analyze the JSON returned by the LLM and all raw LLM responses are available in our project GitHub: github.com/NeuroBridge/LLM-Metadata-Extraction-Paper-2025.

